# Cluster analysis on high dimensional RNA-seq data with applications to cancer research - An evaluation study

**DOI:** 10.1101/675041

**Authors:** Linda Vidman, David Källberg, Patrik Rydén

**Author notes:** **Corresponding author:** Linda Vidman, Department of Mathematics and Mathematical Statistics, Umeå University, SE 901 87 Umeå, Sweden.

## Abstract

Clustering of gene expression data is widely used to identify novel subtypes of cancer. Plenty of clustering approaches have been proposed, but there is a lack of knowledge regarding their relative merits and how data characteristics influence the performance. We evaluate how cluster analysis choices affect the performance by studying four publicly available human cancer data sets: breast, brain, kidney and stomach cancer. In particular, we focus on how the sample size, distribution of subtypes and sample heterogeneity affect the performance.

In general, increasing the sample size had limited effect on the clustering performance, e.g. for the breast cancer data similar performance was obtained for *n* = 40 as for *n* = 330. The relative distribution of the subtypes had a noticeable effect on the ability of identifying the disease subtypes and data with heavily skewed distributions turned out to be difficult to cluster. Both the choice of clustering method and selection method affected the ability to identify the subtypes, but the relative performance varied between data sets, making it difficult to rank the approaches. For some data sets, the performance was substantially higher when the clustering was based on data from only one sex compared to data from a mixed population. This suggests that homogeneous data are easier to cluster than heterogeneous data and that clustering males and females individually may be beneficial and increase the chance to detect novel subtypes. It was also observed that the performance often differed substantially between females and males.

The number of samples seems to have a limited effect on the performance while the heterogeneity, at least with respect to sex, is important for the performance. Hence, by analyzing the genders separately, the possible loss caused by having fewer samples could be outweighed by the benefit of a more homogeneous data.

## Introduction

Diseases like cancer can arise from a multitude of genetic and epigenetic changes. Studying gene expression profiles from tumor samples in cancer patients can reveal information about novel cancer subtypes [1, 2]. However, the problem is challenging since the regulatory mechanisms underlying gene expression are complex and expression levels are affected by external environmental factors like diet and the use of drugs, as well as internal factors like gender and age.

Profiling of gene expression is a way of analyzing the activity of genes, and technologies like RNA-sequencing and microarrays have made it possible to look at gene expressions for thousands of genes simultaneously. Signatures in gene expression are widely used in medical research, e.g. in the assessment of breast cancer [3, 4] and for predicting prognosis in colon cancer [5-7] and ovarian cancer [8-10]. A common aim is to detect novel disease groups, which can be used for personalized treatments, but to become a more useful tool in the field of personalized medicine there is a need for evaluation of profiling methods [11].

A popular approach for detecting novel subtypes of a disease is to use cluster analysis, which is an unsupervised approach for finding groups with similar patterns [12]. It has been frequently used for identifying subtypes of cancer by clustering samples (individuals) with similar gene expression patterns [13-15], as well as for finding groups of genes that have similar profiles over samples [16, 17]. Classical algorithms such as hierarchical clustering and k-means clustering are popular choices, but there are several alternative clustering methods, e.g. self-organizing map (SOM) [18], Partitioning Around Medoids (PAM) [19] and Density-Based Spatial Clustering of Applications with Noise (DBSCAN) [20]. For an extensive overview of available clustering methods, see [21].

The high dimensionality of gene expression data makes detection of novel subtypes a difficult task, which is also complicated by the noisy character of the data [22]. It is therefore common to apply either variable selection or feature extraction prior to the analysis to lower the dimensionality of the problem. A wide range of variable selection and feature extracting methods have been proposed making the number of clustering approaches, defined by the dimensionality reduction method and the clustering algorithm, very large.

The clustering algorithms use different strategies to define groups and can therefore cluster samples in rather dissimilar partitions. Consequently, the relative performance of different clustering methods can be expected to vary among different data sets. Understanding which factors affect the clustering performance is essential in order to select a suitable clustering approach for a specific problem. However, for problems aiming at detecting novel disease subtypes using gene expression data, there are few studies that have evaluated the performance of different clustering approaches. Jaskowiak et al. [23] compared 4 clustering methods and 12 distance measures and concluded that k-medoids and hierarchical clustering with average linkage were in general superior over hierarchical clustering with complete or single linkage. In an article by de Souto et al. [24], 7 clustering algorithms were compared on 35 gene expression data sets; the best result was achieved with a parametric approach assuming a mixture of multivariate Gaussians. Freyhult et al. [25] showed that k-means and hierarchical clustering using Ward’s linkage performed significantly better than PAM and SOM, but they also pinpointed that pre-processing steps could have a major influence on the performance. To our knowledge, limited efforts have been made in order to understand how the sample size and the distribution of the samples affects the clustering performance in high dimensional gene expression data.

A challenge in detecting novel disease subtypes using cluster analysis is that the individuals in the study are often heterogeneous and that they can be grouped with respect to several factors, e.g. disease subtype, age, and gender. The fact that several of the factors may have a pronounced effect on the gene expression reduces the chance that the resulting cluster partition will agree with the partition defined by the disease subtype. We argue that the more heterogeneous a patient group is, the more difficult it will be to achieve a partition that corresponds to the disease subtype.

One solution would be to base the clustering on a relatively homogeneous subset of patients. A possible drawback with this approach may be that including fewer patients will in itself have a negative effect on clustering performance.

In this paper we compare the performance of 30 clustering approaches, defined by which clustering algorithm is used and how the variable selection and feature extraction are performed. The approaches are used to analyze gene expression data (RNA-seq data) from four human cancer tumor types, each with two known subtypes. In particular, we study how the clustering is affected by the characteristics of the data, including the sample size, the skewness of the subtypes and the homogeneity of the individuals.

## Methods

### Data sets

We included four human cancer data sets from The Cancer Genome Atlas (TCGA) network, where the gene expression data were obtained using Illumina HiSeq 2000 RNA Sequencing Version 2. Clinical data together with Level 3 mRNA-seq RSEM data [26] were downloaded from Broad institute GDAC FireBrowse Version: 1.1.35.

Only samples from primary solid tumors were included in the study. For all data sets, the individuals belong to one of two predefined subclasses, see Table 1. We consider these subclasses as the gold standard partitions but it should be noted that there exist several ways of dividing the observations into subclasses.

**Table 1.**
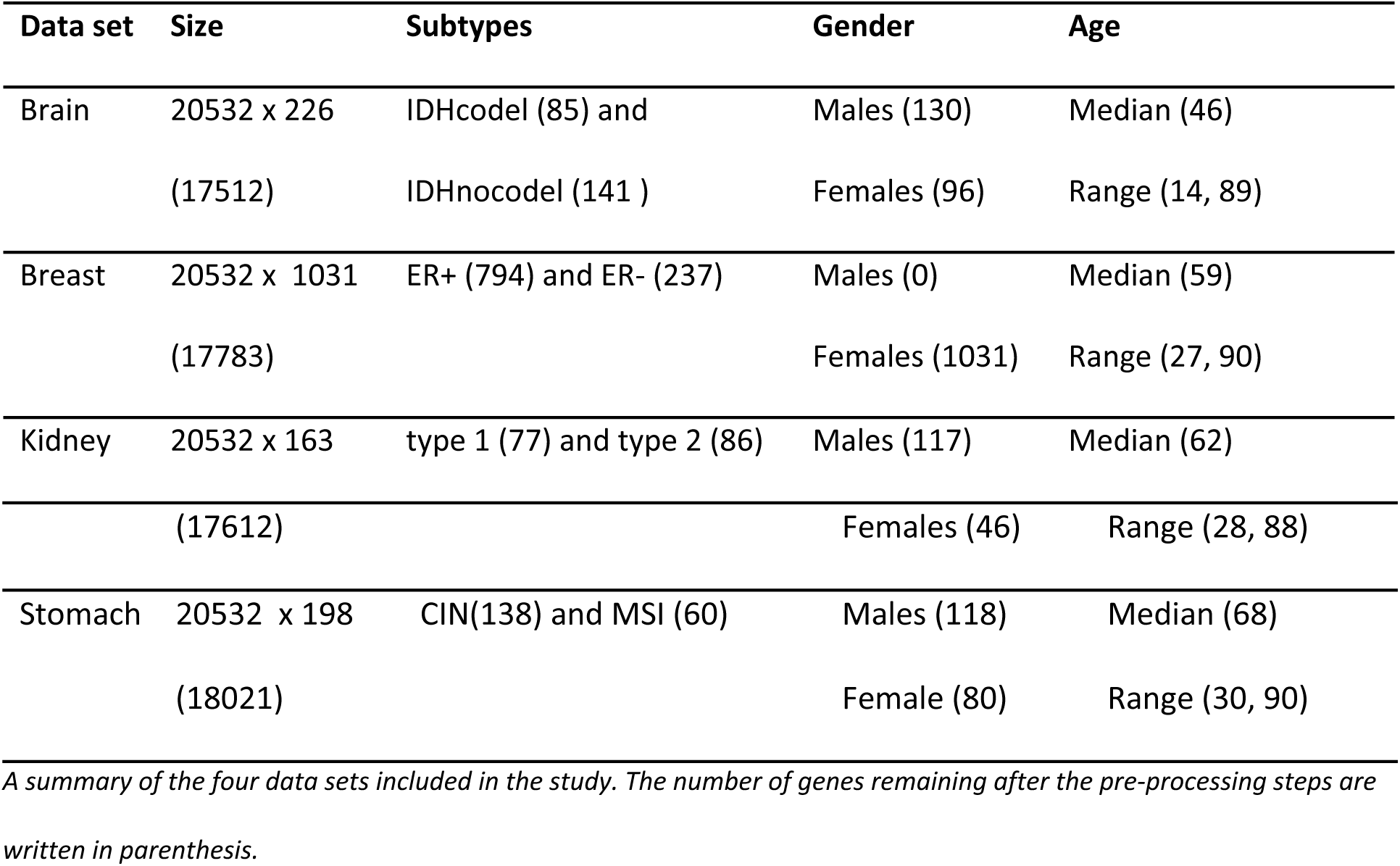
Summary of data sets.

#### Brain cancer data set

The *Brain data* consists of normalized gene expression measurements in tumor samples from patients with lower grade glioma (denoted as LGG by TCGA). The patients were classified into three prognostically significant subtypes, IDHcodel - gliomas with IDH mutation and 1p/19q co-deletion, IDHnocodel-gliomas with an IDH mutation and no 1p/19q co-deletion, and gliomas with wild-type IDH -IDHwt. [27]. Gliomas with wild-type IDH were excluded from the analysis, whereas the two remaining classes defined the subgroups. A total of 226 samples remained after removing samples that were missing information about subtype, see Table 1 for further details.

#### Breast cancer data set

The *Breast data* consists of normalized gene expression measurements in tumor samples from patients with breast invasive carcinoma (denoted as BRCA by TCGA). Subtypes were defined by the Estrogen Receptor (ER) status, either positive or negative [28]. The ER status was given in the clinical data. A total of 1031 samples remained after removing males and samples without subtype information, see Table 1 for further details.

#### Kidney cancer data set

The *Kidney data* consists of normalized gene expression measurements from tumors in patients with kidney renal papillary cell carcinoma (denoted as KIRP by TCGA). Two main subtypes, type 1 and 2, have been histologically determined [29]. A total of 163 cases remained after removing samples that were missing subtype information, see Table 1 for further details.

#### Stomach cancer data set

The *Stomach data* consists of normalized gene expression measurements from tumors in patients with stomach adenocarcinoma (denoted as STAD by TCGA). Four molecular subgroups have been identified: Epstein-Barr Virus (EBV) – positive, Genomically Stable (GS), MicroSatellite Instability (MSI) and Chromosomal INstability (CIN) [30]. Here we kept only the 198 samples classified as CIN and GS and defined those as the two subtypes, see Table 1 for further details.

### Ethics statement

The data included in this study was accessed from TCGA according to open access guidelines and comes from previously publishes articles where written informed consents were obtained in accordance with the local institutional review boards [27, 29-31]. More information is available at: http://cancergenome.nih.gov/abouttcga/policies/informedconsent.

### Pre-processing of data

The analyses were based on normalized level 3 mRNA-seq gene expression data from TCGA. Prior to the analyses the data were log2-transformed. Furthermore, genes that either showed no variation between samples (i.e. standard deviation equal to 0) or were expressed at very low levels (i.e. > 75% of the samples had gene expression values below the 15th percentile) were removed prior to the downstream analysis.

### Variable selection and feature extraction

The first step in the downstream analysis was to determine which variables or features should be included in the cluster analysis. This was done either by selecting the *N* genes with the highest standard deviation, or by creating new features using principal component analysis (PCA) and let the analysis be based on the *M* first principal components (PC). The number of components *M* was set to 5 or 30 and *N* was set at three levels: 100, 1000, and all genes. Hence five *selections methods* were considered: top 100 genes, top 1000 genes, all genes, first 5 PC and first 30 PC. The principal components were obtained using the R-function “prcomp”.

### Cluster analysis

The final step in the analysis was to cluster tumor samples based on their gene expression profiles with the objective to detect cancer subtypes. Clustering algorithms often demand that the user specifies the number of resulting clusters. Here, the analyses were performed in such way that the clustering resulted in two clusters. The main aim here was not to study the relative performance of different clustering approaches, but to study how data characteristics like sample size, distribution of subtypes and sample heterogeneity influence the clustering performance. We therefore decided to include commonly used clustering algorithms and commonly used methods for variable selection and feature extraction. Descriptions of the clustering algorithms considered in this paper are given below.

#### Hierarchical clustering

There are two strategies in hierarchical clustering; agglomerative and divisive. Here the agglomerative clustering was used. This bottom-up approach starts by treating the individual samples as clusters and then recursively joins them until only one single cluster remains. Hierarchical clustering requires that the user specifies a dissimilarity measure and a method for calculating distances between clusters. In this paper both the Manhattan distance and the absolute Pearson correlation distance (i.e. *d*(*x,y*) =1 − | *ρ*(*x,y*) |, where *ρ* is Pearson correlation) were used as dissimilarity measures and Ward’s linkage was used to calculate the distance between clusters [32]. The resulting dendrogram was cut so that two clusters were defined. The R-function “hclust” was used for performing hierarchical clustering. Henceforth the hierarchical cluster analysis method utilizing the Manhattan distance and the absolute Pearson correlation distance will be denoted by hclust(M) and hclust(cor), respectively.

#### K-means clustering

The k-means algorithm (kmeans) partitions *n* samples into *k* clusters where each sample is assigned to the cluster with the nearest mean. The method requires the user to predefine the number of clusters *k,* here *k* = 2 The aim of the algorithm is to minimize the within-cluster sum of squares, *i.e.*

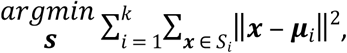

where {***x***_1_, …, ***x***_*n*_} are the d-dimensional observations, ***S*** = {*S*_1_, …, *S*_*k*_} are the *k* clusters and {**μ**_1_, …, **μ**_*k*_} are the associated centroids [33].

The analyses were made using the R-function “kmeans”, which used 10 random starts (nstart = 10).

#### Self-organizing map

Self-Organizing Map (SOM) is a type of artificial neural network that can be used for clustering. A SOM consists of two layers; one input and one output layer. The output layer typically consists of two-dimensional arrays of nodes that are ordered as a rectangular or hexagonal grid. The input layer is fully connected to the output layer, and each node has a vector of weights of the same dimension as the input vector. The weights are updated during the training process and when the weights are fully trained, the distance from each node to the input data is calculated and the sample falls into the cluster (node) that is closest [18].

The clusters were obtained using the function “som” in the R-package “kohonen” [34], with default settings, a hexagonal 2×1 grid and rlen = 1000. Since the function uses a random input, the algorithm was run ten times where only the result which gave the lowest within-variation was reported.

#### Affinity propagation

The Affinity Propagation (AP) algorithm tries to identify data points (exemplars) that can represent clusters. It considers every data point as a potential exemplar and tries to maximize the total similarity between data points and their exemplars. If we consider data points as nodes in a network, then the algorithm operates by sending two types of messages along the edges: one that represent how well suited a data point *x* is to serve as exemplar to a data point *y*, and one indicating how suitable it is for *y* to choose *x* as it’s exemplar. The algorithm updates the messages until convergence [35].

The R-function “apclusterK” in the package “apcluster” [36] with parameters prc = 0, lam = 0.9, maxits = 2000, convits = 200 and nonoise = TRUE was used for performing affinity propagation. The number of clusters *k* was set to 2 and the function “negDistMat” was used for calculating dissimilarities.

#### Cluster ensemble

The resulting clusters from the above mentioned methods were used to create a consensus cluster using Cluster Ensemble (CE). This was achieved using the R-function “cl_consensus” in package “clue” [37], with method “HE”, which is a fixed-point algorithm for obtaining hard Euclidean least squares consensus partitions. The analysis resulted in a consensus partition dividing the samples into two groups.

### Data composition

Aside from the fact that the data sets in this study come from different cancer diseases, there are a number of other things that characterize them as well, e.g. the sample size, the relative distribution of the two subtypes and underlying factors like gender and age distribution.

In order to study how the data characteristics affected the clustering performance, we constructed sub-data sets by sampling the data as described below. Unless otherwise stated the sampling was made without replacement and repeated 10 times to reduce sampling effects. We sampled a maximum of 70% of the available samples. All cluster analysis approaches were applied to each of the sub-data sets.

#### Number of samples

The number of samples in the data sets varied between 163 and 1031. To evaluate how sensitive the clustering performance was to sample size, we compared the performance by considering sub-data sets with different number of samples. The sample size was varied (with a step of 10) from 40-110, 40-330, 40-100 and 40-80 for Brain, Breast, Kidney and Stomach, respectively. Each sampling was made so that the fraction of observations belonging to each of the two subtypes was fixed at 50% (symmetric distribution). In addition, the analysis were repeated for a skewer distribution of subtypes with 20% of the samples in the smallest group. The maximum number of samples were then 120, 270, 70 and 120 for Brain, Breast, Kidney and Stomach respectively.

#### Distribution of subtypes

Here the fraction of samples belonging to the first subtype was varied while keeping the sample size fixed. The percentage of samples belonging to the first subtype was altered at the levels 10%, 20%, 30%, 40% and 50%, with 108, 100, 65 and 83 samples for Brain, Breast, Kidney and Stomach, respectively.

#### Homogeneity with respect to gender

Homogeneous data sets with respect to gender (i.e. data sets including only males or females) were constructed such that the distribution of the subtypes was symmetric and the sample size was the same for both genders. As a control, we constructed a heterogeneous data set with 50% female and 50% male, having the same sample size and relative distribution of the subtypes as the two homogeneous data sets. Here the sample sizes were 55, 28 and 36 for Brain, Kidney and Stomach, respectively. The ARIs for the heterogeneous data were calculated separately for males and females.

### Difficulty of the clustering problems

The considered sub-classes for the data sets were regarded as gold standards. Arguably, other divisions of the data could have been considered. A minimum criterion on a gold standard is that there should be a genetic signal that can be detected using supervised teckniques. To assess how well the gene expression data separated the cancer subtypes, we used PCA and plotted the first two components with the patients colored according to subtypes. In addition, the supervised classifier random forest was applied to the data sets with the gold standard partition as response [38]. The fraction of correctly classified samples was used as a measure of the separation between the subtypes. A data set with low separation may be regarded as a difficult clustering problem. Random forest using sampling with replacement was performed using the R-package “randomForest” [39].

### Performance measure

The performance of a clustering approach was determined by how well the resulting partition matched the gold standard partition. Several clustering similarity measures have been proposed [40-42]. We used the adjusted Rand index (ARI), which is a modified version of the Rand index (proposed by William Rand 1971) that adjusts for agreement by chance [43].

ARI is calculated as follows. Suppose we have *n* objects in a set *S*. Let *A* = {*A*_1_, *A*_2_, …, *A*_*l*_} and *B* = {*B*_1_, *B*_2_, …, *B*_*m*_} be two partitions of *S* (one is the gold standard partition and one is the result from clustering). Then the overlap of *A* and *B* can be described by the following contingency table

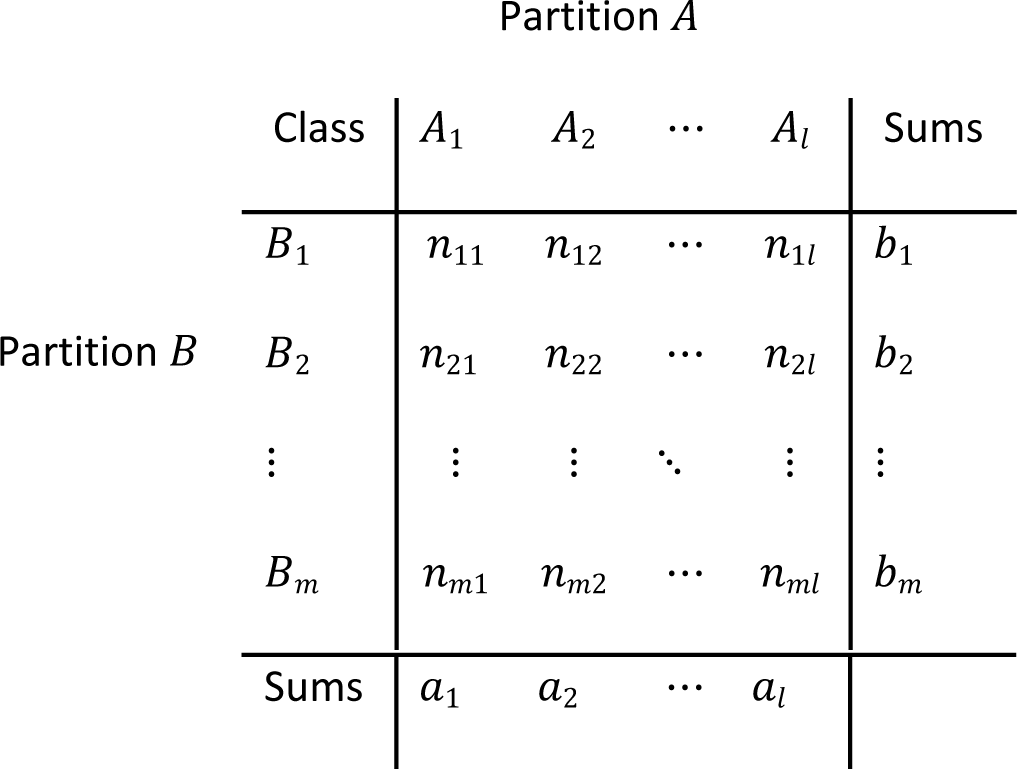

where *n*_*ij*_ denotes the count of common objects in *A*_*j*_ and *B*_*i*_.

The Adjusted Rand Index (ARI) is then obtained as

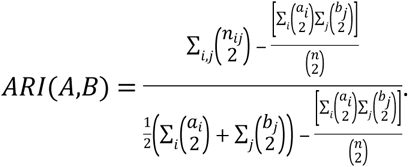

The ARI takes value 1 if the two partitions are identical and has expected value 0 in the case of a completely random partition. It can yield negative values if the agreement is worse than expected by chance. The R-function “RRand” in package “phyclust” [44] was used for calculation of ARI.

The same ARI-value can be obtained by very different partitions of the samples. Hence the fact that two cluster outputs yield the same ARI-values compared to the gold standard does not necessarily imply that the resulting clusters are similar. In addition to quantifying the agreement with the gold standard, ARI was used to quantify the similarity between different clustering approaches, by considering the *average of the pairwise ARI* (apARI).

The agreement between partitions was also evaluated using the distance metric *variation of information* [40]. Since the relative results were very similar to those obtained by ARI, we chose to present only the ARI-values.

### Statistical analyses

In the cases where we have subsampled data, the tables contain mean values of the 10 replicates. For comparisons between different settings (e.g. *n* = 40 versus *n* = 100) we used the one-sample Wilcoxon signed rank test based on the values from the 30 clustering approaches given in the tables (clustering approaches using 30 PC were omitted for comparisons with sample size lower than 30). The Wilcoxon rank sum test was used in the case when we compared apARI for different sample sizes. For each test we report the median of the ARI-differences (i.e. delta ARI) and the corresponding p-value (p). Note that the figures were obtained in a slightly different way than the tables. The figures show the result for the 10 replicates, where each value was obtained by averaging over all 30 clustering approaches. All analyses were performed using the R programming language version 3.4.3 [45].

## Results

All data included in this study were derived from tumor samples in cancer patients with known subtypes. The partition defined by the subtypes served as the gold standard. Brain and Breast revealed clear separation between the subtypes, see Fig 1. Supervised classification yielded 99.1% and 91.7% accuracy for Brain and Breast, respectively. The subtypes were less distinguishable for Stomach and Kidney, with a classification accuracy of 73.6% and 86.9%, respectively, see Fig 1. Hence, all data sets contain information that distinguish the subtypes although Brain and Breast appear to be relatively easier clustering problems.

**Fig 1.**
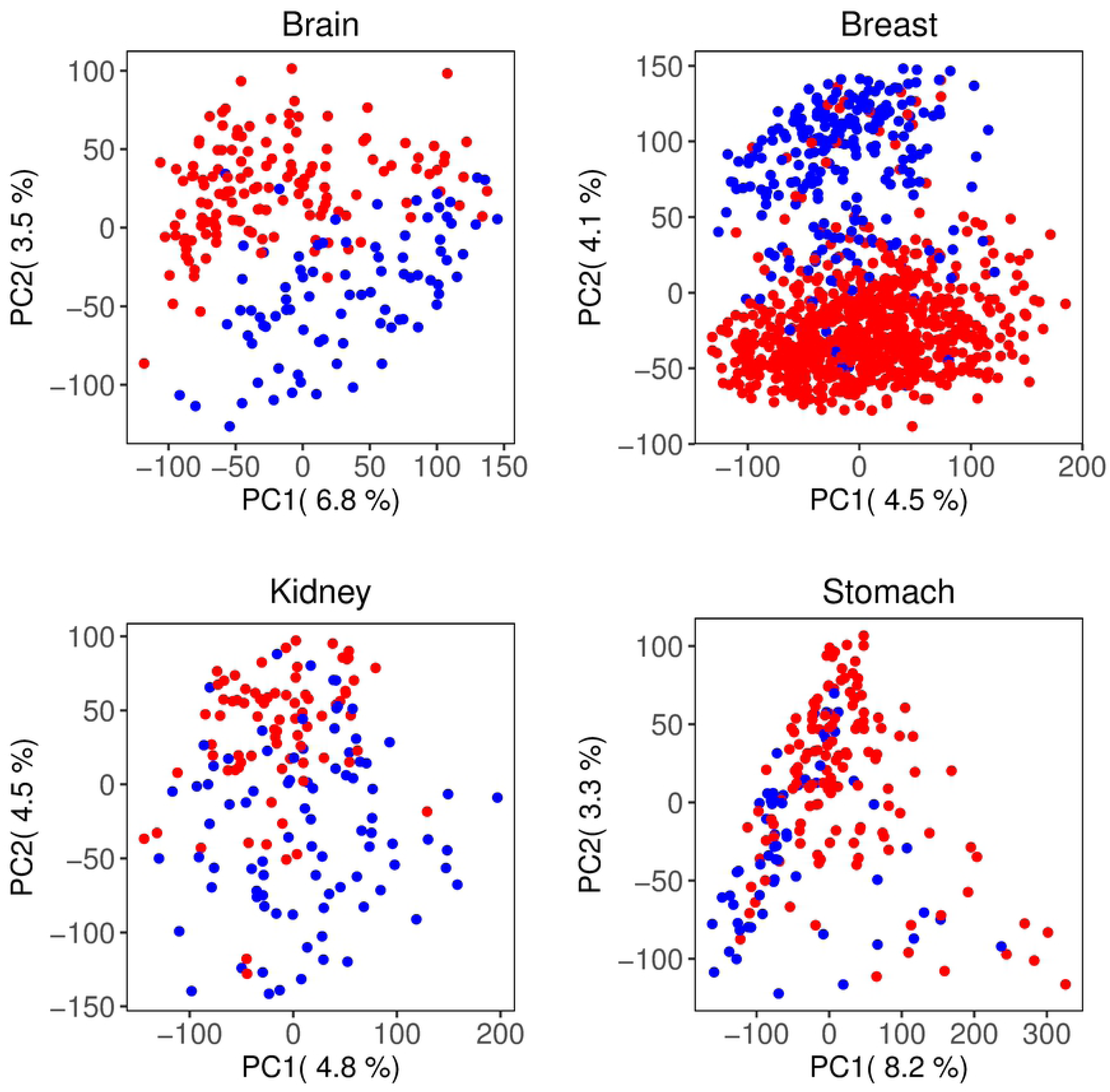
Principal component analysis to visualize separation between subtypes. Figures based on the first two principal components, where the subtypes are marked in different colors; Breast: ER+(red) and ER-(blue), Brain: IDHnocodel (red) and IDHcodel (blue), Kidney: type 1(red) and type 2(blue), Stomach: CIN (red) and MSI (blue).

Here we present results on how clustering performance was affected by clustering method, selection method, sample size, relative distribution of subtypes and the homogeneity of the data.

### Clustering approaches

Thirty clustering approaches (six clustering methods combined with five selection methods) were applied to the four data sets, where the performance was quantified using ARI. The clustering performance varied considerably between the data sets. Breast was relatively easy to cluster while the other data sets were harder to cluster. The median ARI (taken over all 30 clustering approaches) for Brain, Breast, Kidney and Stomach were 0.10, 0.55, 0.17 and 0.15 respectively. Interestingly, the overall clustering performance for Brain was low while the obtained accuracy using supervised classification was relatively high. This shows that a simple classification problem is not necessarily an easy clustering problem.

The choice of clustering method had a large impact on the clustering performance, but the methods’ relative performance varied between the data sets making it difficult to rank the methods, see Fig 2A and S1 Table. Furthermore, the performance was often sensitive to the choice of selection method, e.g. when applying SOM to Breast the ARI ranged from 0.00 (5 PC) to 0.69 (30 PC).

**Fig 2.**
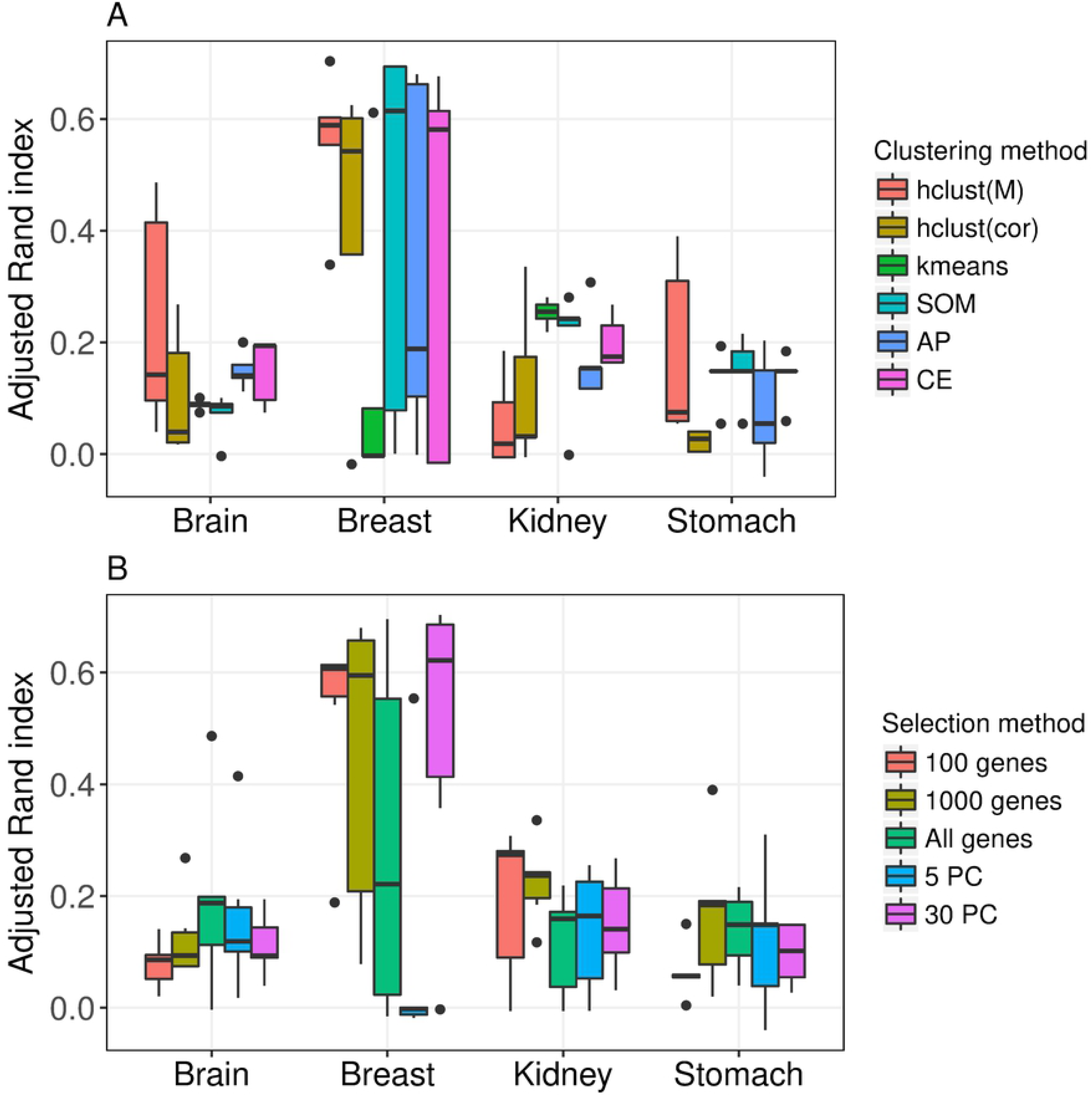
Performance of clustering methods and selection methods. Adjusted Rand index for clustering result compared to gold standard partition. Figure A shows results for the clustering methods, where each box contains observations from the five selection methods. Figure B shows results for the selection methods, where each box contains observations from the six clustering methods.

The relative performance of the five selection methods also varied between the data sets, making it hard to draw general conclusions, see Fig 2B and S1 Table. However, two clustering approaches, hclust(cor) combined with 5 PC or 30 PC, had relatively low performance for all data sets, see S1 Table.

### Sample size

Next we investigated how the sample size affected the performance of the clustering, while keeping the distribution of subtypes fixed at 50%. To our surprise, only limited gain in performance was observed when the sample size was increased, see Fig 3 and S2-S5 Table. In Brain, a small significant increase in performance (delta ARI = 0.016, p = 0.014) was observed between the smallest and largest considered sample sizes (*n* = 40 and *n* = 110). For Stomach we found no significant increase in performance (delta ARI = −0.004, p = 0.524) between the smallest (*n* = 40) and largest (*n* = 80) sample sizes. This may partially be explained by the fact that the performance was very low also when the sample size was relatively high, see Fig 3. For Kidney, there was a modest increase in performance when *n* was increased from 40 to 100 (delta ARI = 0.091, p <0.0001), but there was no stable trend, see Fig 3. For Breast, no significant increase in performance was found between *n* = 40 and *n* = 330, (delta ARI = −0.028, p = 0.529). The trends presented were observed independently on which clustering algorithm was considered, see S2-S5 Table.

**Fig 3.**
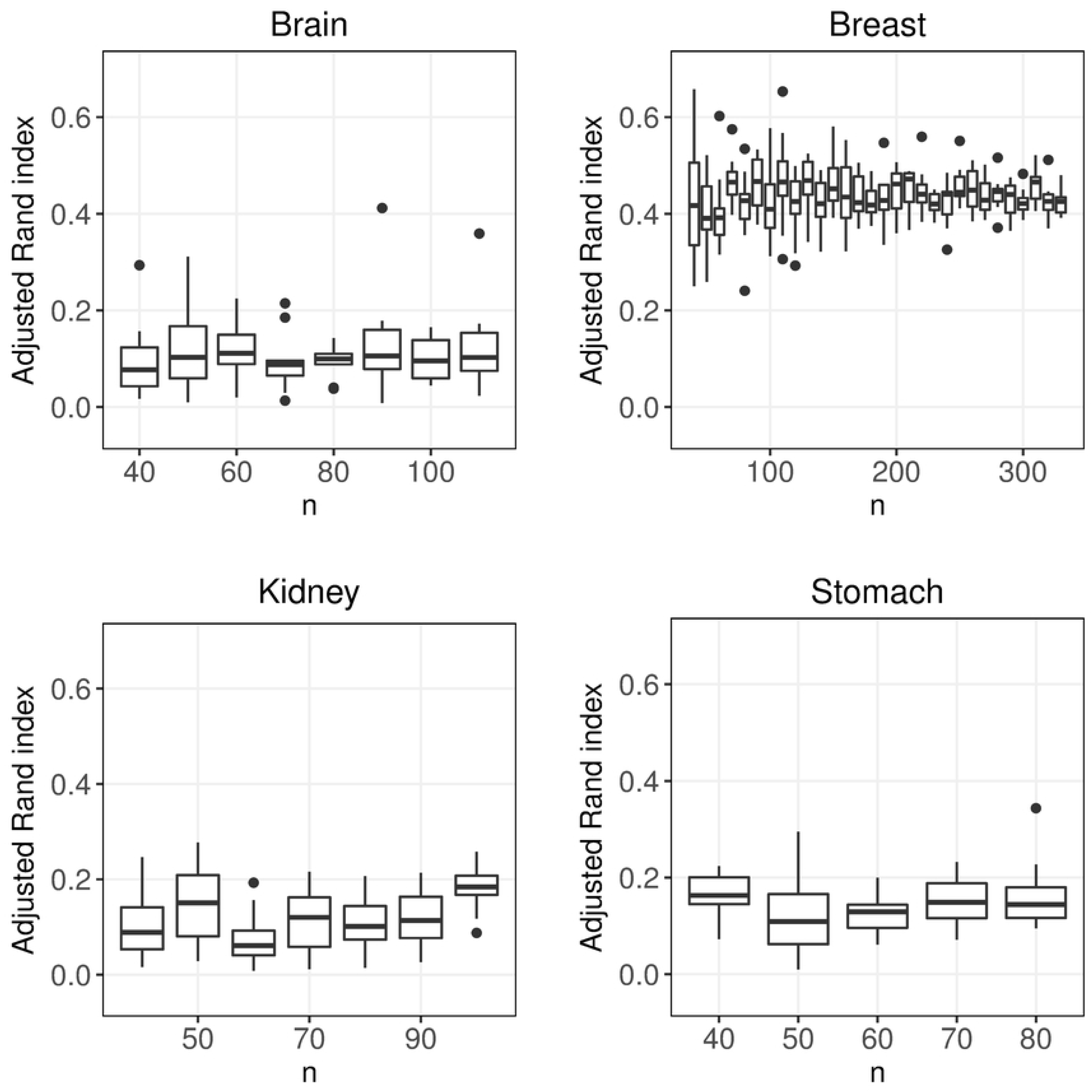
Sample size. Boxplots of adjusted Rand index for different number of observations and a symmetric distribution of the subtypes. Each box contains mean adjusted Rand index values (taken over of all 30 clustering approaches) for 10 replicates.

In addition to consider the symmetric distribution, with 50 % of each of the subtypes, we also considered a case with a more skewed distribution (20% of the observation in the smallest group). Again, it was observed that the sample size had a limited effect on the performance, see S6-S9 Table.

Interestingly, although increased sample size had limited effect on the performance it was evident that the 30 approaches clustered the individuals more similar when the sample size increased from relatively small (*n* = 40) to modest (*n* = 100), see Fig 4. This was most evident for Breast, where the average pairwise ARI (apARI) was significantly lower at *n* = 40 than at *n* = 330 (delta apARI = 0.204, p <0.001). For all data sets, the agreement between the approaches was higher than the agreement with the gold standards. Arguably, this indicates that there are relatively strong signals in the data sets that are not in absolute agreement with the considered cancer subtypes, see Fig 3 and 4.

**Fig 4.**
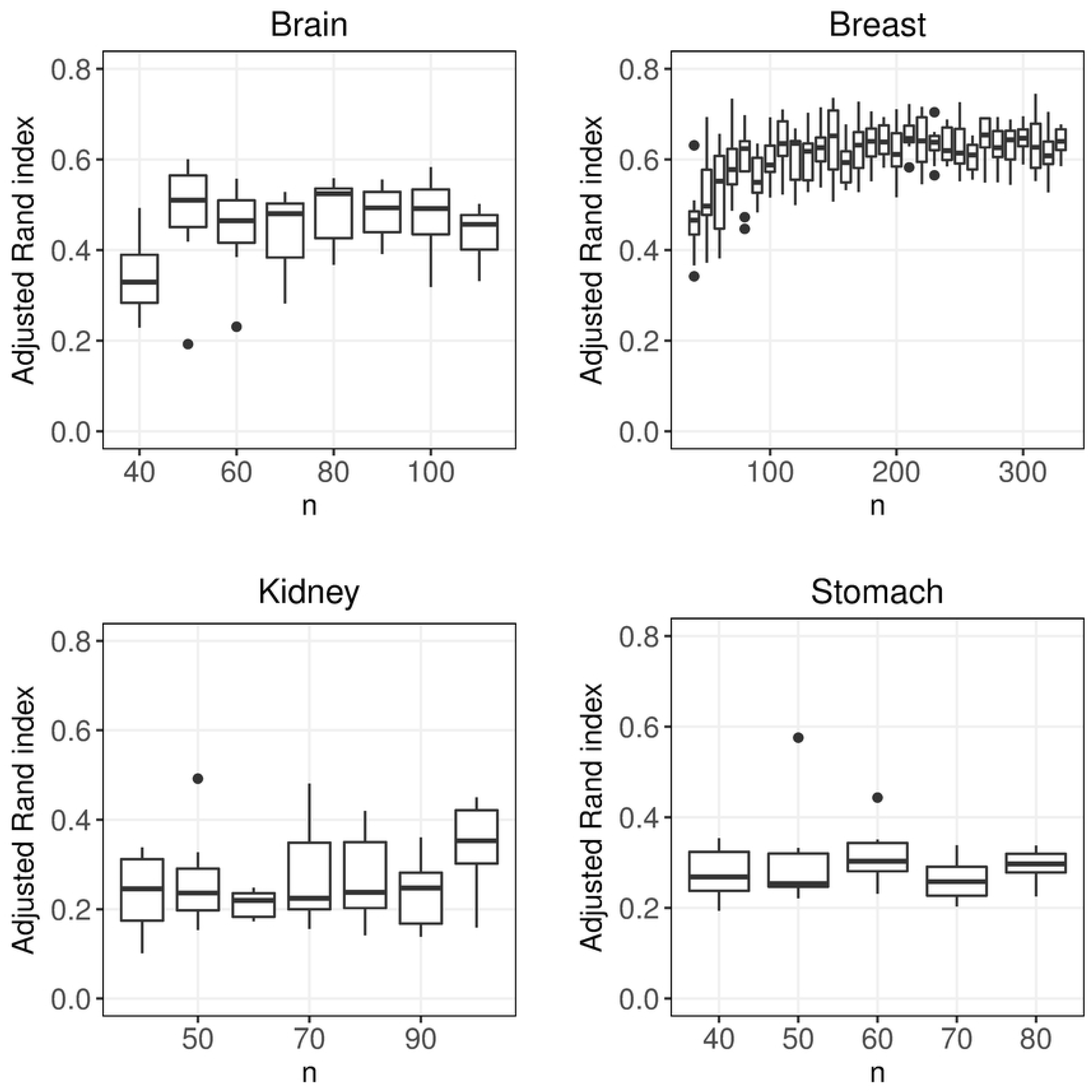
Similarity between clustering approaches for different sample sizes. Average pairwise adjusted Rand index (apARI) between clustering approaches for different number of observations.

### Distribution of subtypes

We investigated how the performance was affected when the relative distribution of the two subtypes was altered. Here the sample size was fixed while the proportion of the smallest subtype was altered from 10% (skewed data) to 50% (symmetric data). It should be stressed that the relative distribution of the classes in practice are unknown, and cannot be estimated in advance. For Brain, Breast, Kidney and Stomach, the performance was significantly higher for the symmetric data compared to the skewed data, (delta ARI = 0.022, p < 0.001), (delta ARI = 0.481, p <0.0001), (delta ARI = 0.102, p <0.0001) and (delta ARI = 0.106, p <0.0001) respectively, see Fig 5 and Table S10-S13. Somewhat surprisingly, the highest median ARI-values were found at 40/60 distribution (Brain, Kidney and Stomach) and at 30/70 distribution for Breast.

**Fig 5.**
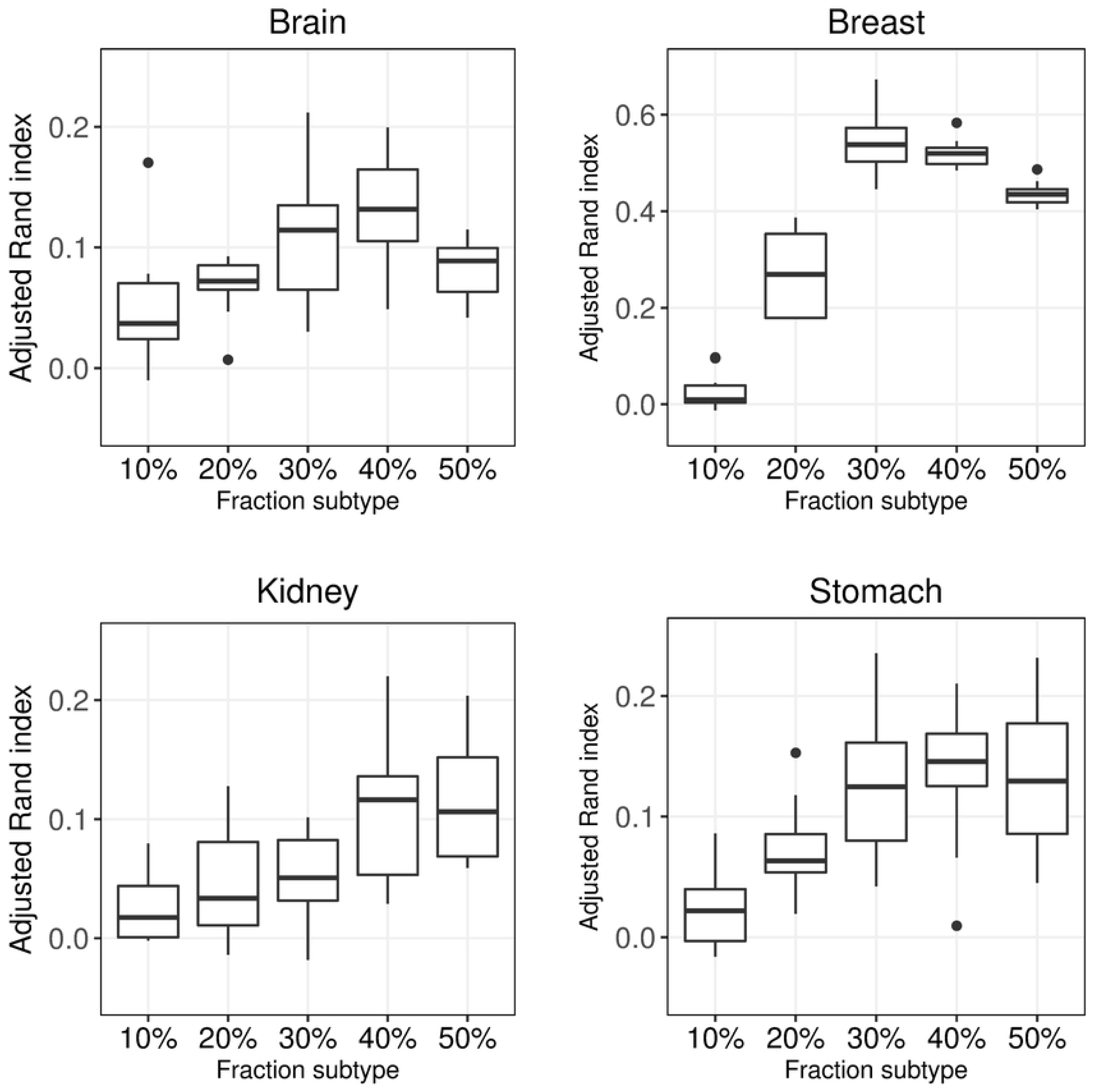
Distribution of subtypes. Adjusted Rand index for subtype fractions 10% – 50%. Each box contains mean adjusted Rand index values (taken over of all 30 clustering approaches) for 10 replicates.

Breast was investigated further by considering subtype fractions between 10-90%, see S11 Table. Here the most skewed data with 10% or 90% of the smallest subtype yielded very low performance with median ARI-values −0.003 and 0.084, respectively. Less skewed data with 20% or 80% performed better, with median ARI-values 0.309 and 0.205, respectively. The fairly symmetric data sets with 30%, 40%, 50%, 60% and 70%, had the best performances with median ARI-values 0.594, 0.552, 0.474, 0.398 and 0.355, respectively. Almost all of the differences between the fractions were significant, the exception was the 20% fraction that did not differ significantly from the 70% fraction.

Investigating the clustering methods with respect to skewness revealed that all clustering methods were affected by the skewness of the data, but for hclust(cor) the differences were less visible due to the overall low performance, see S14 Fig and S10-S13 Table.

### Analyze genders separately or together?

It is well known that there are widespread gender differences in gene expression [46, 47]. Hence, gender may influence gene expression and the clustering result, suggesting that it would be relatively easier to cluster data that are homogeneous w.r.t. sex. Furthermore, there is a possibility that the genomic signal for different subtypes may vary in strength between males and females. To study how gender influenced the clustering, the data sets (Brain, Kidney and Stomach) were used to construct new data sets: only females (Females), only males (Males) and mixed group with 50% females. The sample size and the relative distribution of the subtypes were kept fixed. For the mixed group, clustering was performed on all samples but ARI was calculated in three ways: using all patients (Mixed all), only the females (Mixed females) and only males (Mixed males), see Methods for further details.

Arguably, a relative high ARI for Females (Males) compared to Mixed females (Mixed males) suggests that a homogeneous data w.r.t. sex has a positive effect on the clustering performance. The ARI-values for Females and Males was used to compare the strength of the subtype specific signals.

The clustering performance was higher when the clustering was performed on homogeneous data compared to heterogeneous data in all cases but one, see Fig 6 and S15 Table. The biggest differences were observed for Kidney and Stomach. For Kidney median ARI was 0.049 for Mixed Males and 0.159 for Males (225 percentage increase, delta ARI = 0.092, p <0.0001) the corresponding numbers for females were (84 percentage increase, delta ARI = 0.027, p = 0.042). For Stomach median ARI was 0.095 for Mixed Males and 0.183 for Males (93 percentage increase, delta ARI = 0.091, p <0.0001) the corresponding numbers for females were (20 percentage increase, delta ARI = 0.027, p = 0.032). For Brain the differences between analyses using homogeneous and heterogeneous data were relatively small (delta ARI = 0.033, p = 0.109) and (delta ARI = −0.016, p = 0.001) for males and females respectively. For Brain and Kidney, males were considerably easier to cluster than females (delta ARI = 0.115, p < 0.0001) and (delta ARI = 0.100, p < 0.0001) respectively. For Stomach, no significant difference was observed (delta ARI = −0.002, p = 0.903). An interesting finding is that the above results holds true for almost all of the considered clustering approaches, see S15 Table.

**Fig 6.**
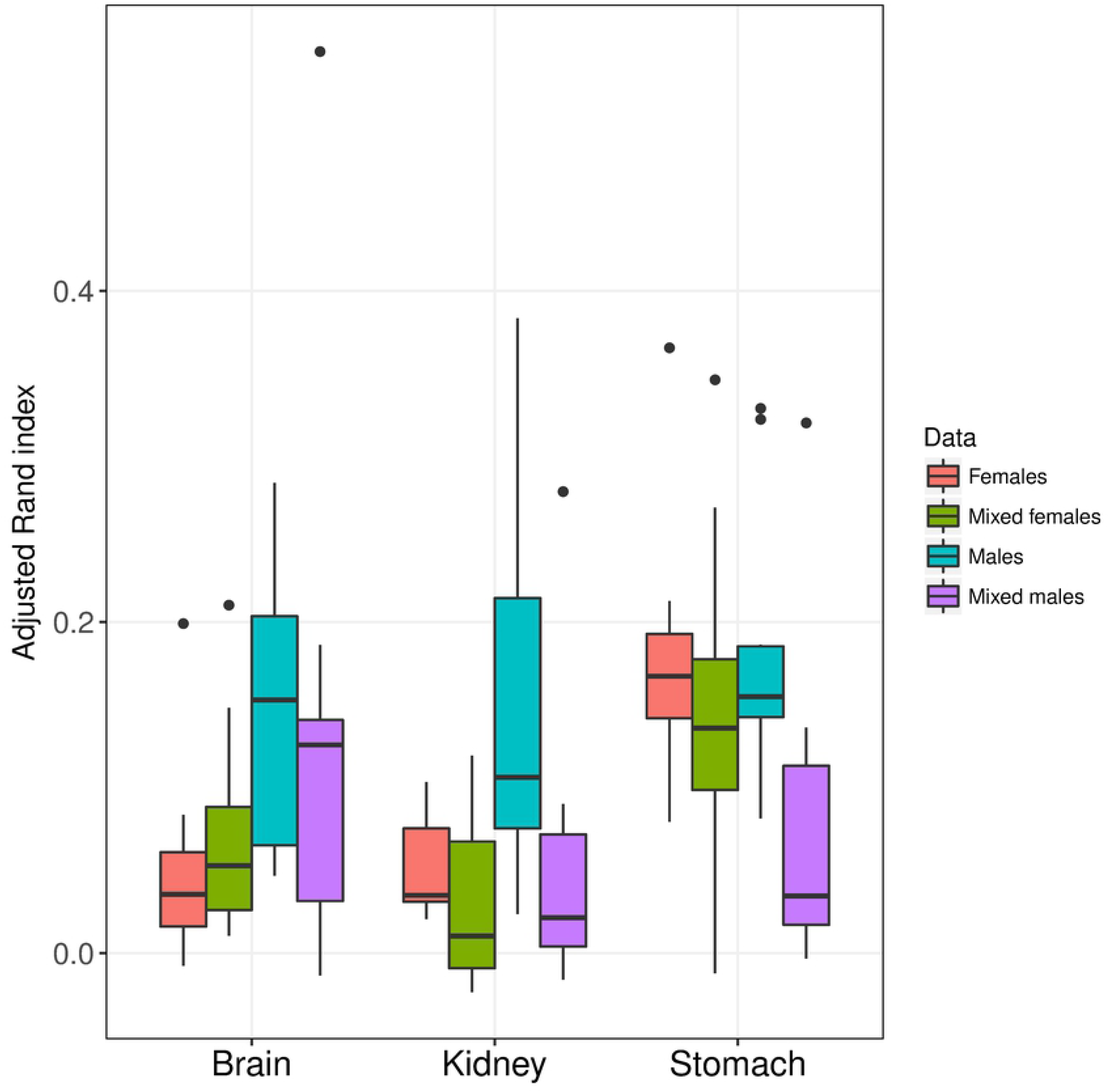
Gender difference. Adjusted Rand index for Brain, Kidney and Stomach when dividing samples by gender. All data sets had a symmetric distribution (i.e. 50 % of each subtype).

## Discussion

The overall aim with this work was to identify and quantify sources that affect the performance of the clustering. Arguably, unsupervised classification, where the objective is to discover novel subgroups, is among the most difficult statistical problems. The two major difficulties being that we are unable to train our model since the true partitions of the samples is not known and that the data generally are affected by several factors and not just the factor of interest, e.g. the factor defining the novel cancer subtypes. For example, patients are diverse with respect to gender, age, ethnicity, etc. and all factors may contribute to differences in gene expression. The lack of a true answer makes it hard to validate and evaluate clustering approaches, which is a serious problem since applying different clustering approaches often yield very different results.

In this work the gold standard partitions were defined by two known subtypes. Although our ambition was to select as relevant subtypes as possible it is clear that there are several alternative partitions that could have been considered for validation. Here, the focus was to examine the relative effect of applying different clustering approaches and to study the impact sample size, skewness and data heterogeneity had on the performance. Arguably, this makes the definition of the gold standard less important.

Supervised classification suggested that there was a strong genetic signal (i.e. the subtypes could be predicted) in all four data sets. Despite this, clustering resulted in relatively low performance for Brain, Kidney and Stomach, although higher than expected by chance. This may suggest that the data are strongly affected by factors that are not necessarily related to the considered subtypes. Arguably, clustering homogeneous data should be easier than clustering heterogeneous data. This was confirmed when the genders were analyzed separately, where the performance was substantially higher when the clustering was based on only one sex compared to a mixed population. Splitting the samples into more homogeneous sets inevitably leads to fewer samples, which may have a negative effect on the clustering performance.

To our initial surprise, increasing the sample size had little influence on the performance. We did get similar performances independently if we used 40 or 300 samples. However, the performances of the clustering approaches became more similar when the sample size increased. Arguably, this suggest that sample size has some importance, but that increasing the sample size per se will not solve the clustering problem. It should be stressed that we did not study how the performance was affected when very small samples (*n* > 20) were considered.

Even though there are recognized gender differences in both cancer survival and treatment response, it is not common that the clustering of subtypes are made separately for males and females [48]. Based on our findings we believe that many omics cluster analysis studies would benefit from analyzing smaller but more homogeneous data sets. In addition, clear gender differences were observed. For Brain and Kidney, the male data were considerably easier to cluster than the female data, suggesting a gender specific genetic signal or that the male group is more homogeneous.

The choice of clustering and selection methods affected performance, but the ranking of the methods varied between the data sets making it difficult to draw general conclusions. However, hclust(cor) often yielded lower performance than the other methods, especially when the selection method involved principal components.

The clustering performance was sensitive to the relative distribution of the subtypes, where data with heavily skewed distributions turned out to be difficult to cluster. The distribution of the subtypes is not known in advance, but if the subtypes are believed to be associated with difference in survival (or some other variable), it may be possible to use survival data to get an idea of the subtype distribution.

## Conclusions

Clustering high-dimensional gene expression data is a challenge. Even with a strong genetic signal, cluster analysis approaches may fail to identify the partition of interest. One important reason for this is that gene expression data are influenced by several factors, which may or may not be known to the researcher. Furthermore, the optimal clustering approach depends on the data and general recommendations are therefore difficult to give. However, the results suggest data characteristics may influence the clustering performance. Interestingly, increasing the sample size may not enhance the performance of the clustering although it make the clustering approaches more similar. If the distribution of the subtypes is skewed, clustering can be very difficult. Finally, the result shows that homogeneous data are easier to cluster than heterogeneous data. Together this suggests that it may be beneficial to analyze the genders separately. The gain of obtaining a more homogeneous data outweighs the possible drawback of having fewer samples.

## Acknowledgements

Not applicable

## Supporting Information

***S1 Table. Performance for different clustering approaches***. Adjusted Rand index for different data sets and 30 clustering approaches (combination of clustering method and selection method). (XLSX)

***S2 Table. Performance for different sample sizes in the Brain data set.*** Average adjusted Rand for different sample sizes (*n*) a 50/50% distribution of subtypes. The table shows the mean values based on 10 random sub-sampled data sets. (XLSX)

***S3 Table. Performance for different sample sizes in the Breast data set.*** Average adjusted Rand for different sample sizes (*n*) and a 50/50% distribution of subtypes. The table shows the mean values based on 10 random sub-sampled data sets. (XLSX)

***S4 Table. Performance for different sample sizes in the Kidney data set.*** Average adjusted Rand for different sample sizes (*n*) a 50/50% distribution of subtypes. The table shows the mean values based on 10 random sub-sampled data sets. (XLSX)

***S5 Table. Performance for different sample sizes in the Stomach data set.*** Average adjusted Rand for different sample sizes (*n*) a 50/50% distribution of subtypes. The table shows the mean values based on 10 random sub-sampled data sets. (XLSX)

***S6 Table. Performance for different sample sizes in the Brain data set.*** Average adjusted Rand for different sample sizes (*n*) for a 20/80% distribution of subtypes. The table shows the mean values based on 10 random sub-sampled data sets. (XLSX)

***S7 Table. Performance for different sample sizes in the Breast data set.*** Average adjusted Rand for different sample sizes (*n*) for a 20/80% distribution of subtypes. The table shows the mean values based on 10 random sub-sampled data sets. (XLSX)

***S8 Table. Performance for different sample sizes in the Kidney data set.*** Average adjusted Rand for different sample sizes (*n*) for a 20/80% distribution of subtypes. The table shows the mean values based on 10 random sub-sampled data sets. (XLSX)

***S9 Table. Performance for different sample sizes in the Stomach data set.*** Average adjusted Rand for different sample sizes (*n*) for a 20/80% distribution of subtypes. The table shows the mean values based on 10 random sub-sampled data sets. (XLSX)

***S10 Table. Performance for different subtype distributions in Brain data.*** Average adjusted Rand index for 10 random samplings for different proportions of subtypes in the Brain data set. (XLSX)

***S11 Table. Performance for different subtype distributions in Breast data.*** Average adjusted Rand index for 10 random samplings for different proportions of subtypes in the Breast data set. (XLSX)

***S12 Table. Performance for different subtype distributions in Kidney data.*** Average adjusted Rand index for 10 random samplings for different proportions of subtypes in the Kidney data set. (XLSX)

***S13 Table. Performance for different subtype distributions in Stomach data.*** Average adjusted Rand index for 10 random samplings for different proportions of subtypes in the Stomach data set. (XLSX)

***S14 Fig. Performance for different subtype distributions and clustering methods.*** Adjusted Rand index for 10 random samplings and 5 gene selection methods. (TIF)

***S15 Table. Performance for genders with even subtype distribution***. Average adjusted Rand index for 10 random samplings when keeping the subtypes fixed at 50%. (XLSX)

